# Changes in nuclear and actin mechanics from G1 to G2 affect nuclear integrity

**DOI:** 10.1101/2025.04.16.649171

**Authors:** Samantha Bunner, Katie Huang, Anish Shah, Nick Lang, Catherine Chu, Schuyler Figueroa, Nebiyat Eskndir, Mai Pho, Gianna Manning, Lilian Fritz-Laylin, Katrina B Velle, Joshua Marcus, James Orth, Andrew D. Stephens

## Abstract

The structural integrity of the nucleus is dependent on nuclear mechanical elements of chromatin and lamins to resist antagonistic actin cytoskeleton forces. Imbalance results in nuclear blebbing, rupture, and cellular dysfunction found in many human diseases. We used Fluorescent Ubiquitin Cell Cycle Indicator (FUCCI) cells to determine how cell cycle changes affect the nucleus and actin force balance. While nuclear blebs are present equally throughout interphase, nuclear blebs form predominantly in G1 and then persist into G2 due to increased actin-based nuclear confinement and focal adhesion density in G1 vs. G2 cells. Upon artificial confinement, G2 nuclei ruptured more than G1 nuclei. Single nucleus micromanipulation force measurements confirmed that G1 nuclei are stronger than G2 nuclei in both the chromatin-based and lamin-based nuclear stiffness regimes. Decreased nuclear stiffness can be explained by loss of peripheral H3K9me3 from G1 to G2, recapitulated by H3K9me3 inhibition via Chaetocin. Cell cycle-based changes in nuclear and actin mechanics impact nuclear integrity and shape.

## Introduction

The nucleus is the organelle that houses the genome and its essential functions. Nuclear mechanics, shape, and integrity are the physical properties that ensure proper nuclear function. Loss of these physical properties is well-documented across the human disease spectrum including heart disease, aging, and cancer (Stephens *et al*., 2019a; Kalukula *et al*., 2022). Nuclear blebs are a specific type of nuclear deformation > 1 µm in size hallmarked by decreased DNA density (Bunner *et al*., 2024; Chu *et al*., 2025; Pujadas Liwag *et al*., 2025). These highly curved nuclear blebs have a high propensity to result in nuclear rupture (Stephens *et al*., 2018; Xia *et al*., 2018). Loss of nuclear mechanics leading to nuclear blebbing and rupture causes nuclear dysfunction via increased DNA damage (Denais *et al*., 2016; Irianto *et al*., 2016; Raab *et al*., 2016; Chen *et al*., 2018; Xia *et al*., 2018; Stephens *et al*., 2019b; Earle *et al*., 2020; Nader *et al*., 2021; Shah *et al*., 2021; Pho *et al*., 2023), changes in transcription (De Vos *et al*., 2011; Helfand *et al*., 2012; Berg *et al*., 2023), and perturbations to cell cycle control (Pfeifer *et al*., 2018) all believed to aid disease progression. It remains unknown how changes during interphase stages impacts nuclear blebbing and rupture.

Changes during the cell cycle could affect nuclear and actin mechanics. For example, genome replication and the resulting doubling of genome content could cause alterations to chromatin, a major mechanical component of the nucleus. More specifically, chromatin histone modifications (Shimamoto *et al*., 2017; Stephens *et al*., 2017, 2018, 2019b; Hobson *et al*., 2020; Nava *et al*., 2020; Williams *et al*., 2020; Danielsson *et al*., 2022; Manning *et al*., 2025), H1 dynamics (Furusawa *et al*., 2015; Senigagliesi *et al*., 2019), chromatin linking proteins (Strom *et al*., 2021; Williams *et al*., 2024) and chromosome conformation interactions (Belaghzal *et al*., 2021) are all known contributors to nuclear mechanics and/or shape that might be altered during or after genome replication. DNA transcription activity has been shown to be essential for nuclear blebbing through a non-bulk mechanical property hypothesized to be due to chromatin motion (Berg *et al*., 2023). Lamins, the other major mechanical component of the nucleus (Dahl *et al*., 2004; Lammerding *et al*., 2006; Swift *et al*., 2013; Stephens *et al*., 2017; Hobson *et al*., 2020; Vahabikashi *et al*., 2022), are known to be added to the nucleus as it grows in size during replication and throughout interphase. Finally, actin confinement and contraction are known to be the antagonistic forces that cause nuclear deformations and rupturing (Le Berre *et al*., 2012; Hatch and Hetzer, 2016; Cho *et al*., 2019; Mistriotis *et al*., 2019; Nmezi *et al*., 2019; Xia *et al*., 2019; Pho *et al*., 2023). Specific changes in actin have been reported at the end of interphase due to decreased focal adhesions (Jones *et al*., 2018), which directly affect the actin cap (Kim *et al*., 2012) which confines and antagonizes nucleus and its shape (Pho *et al*., 2023). These changes are more relevant in constant cycling cells, for example cancer cells, that more frequently exist in S or G2 compared to most cells which remain in G1 or G0 state.

The interphase portion of the cell cycle includes three stages. Gap 1 (G1) is the majority of the interphase cell cycle in which the nucleus exists as a diploid genome. In Synthesis (S) the genome is replicated by DNA polymerase and its associated proteins. The nucleus grows during this stage that is independent of replication (Iida *et al*., 2022). S phase is followed by Gap 2 (G2) in which the nucleus continues moderate growth for a short period of time and prepares to enter mitosis. Tracking cells through the cell cycle can be achieved through use of Fluorescent ubiquitination-based cell cycle indicator (FUCCI) cell lines that have fluorescently labeled Cdt1 and Geminin which respectively are expressed during G1/S and S/G2 (Sakaue-Sawano *et al*., 2008; Marcus *et al*., 2015). Furthermore, cells actively undergoing replication can be labeled via BrdU a DNA analog (Gratzner *et al*., 1975). Nuclear size also correlates with interphase stage. Established cell cycle inhibitors that stall cells in G1 via lovastatin (Rao *et al*., 1999) and G2 via RO-3366 (Vassilev *et al*., 2006) provide a means to directly modulate the interphase stages.

Using different interphase stage indicators, we measured how nuclear shape, integrity, and mechanics are affected by the different stages of interphase. Population images revealed that nuclear blebs are present at levels similar to total population distributions. Timelapse imaging reveal that blebs formed predominantly in G1 and persisted into other interphase stages. Imaging nuclear heights revealed that G1 nuclei are under greater actin confinement (shorter nuclear height) and had more focal adhesions than G2 nuclei. When G1 and late S/G2 nuclei are placed under similar confinement using an artificial confiner, late S/G2 nuclei displayed more frequent ruptures than G1 nuclei. Finally, dual micropipette micromanipulation single nucleus force measurements confirm that G1 nuclei are stronger than G2 nuclei. Immunofluorescence measurements of histone modification states and lamin levels were performed to determine the basis for this nuclear stiffness change. We provide novel findings that both nuclear and actin mechanics change throughout interphase to affect nuclear blebbing and rupture which are applicable to human disease.

## Results

### Nuclear blebs are present equally across interphase cell cycle stages

A nuclear bleb is a protrusion >1 um of the nucleus that forms during interphase and is hallmarked by loss DNA density relative to the main nuclear body (Bunner *et al*., 2024; Chu *et al*., 2025; Pujadas Liwag *et al*., 2025).To determine if nuclear blebs were overrepresented in a particular stage of the interphase portion of the cell cycle, we population imaged static wild type and perturbed HT 1080 Fluorescent ubiquitination-based cell cycle indicator (FUCCI) human cells (Marcus *et al*., 2015). Using FUCCI cells we can measure interphase stage via fluorescent mKO-Cdt1 which increases in G1 and then is degraded gradually upon entering S phase while mAG-Geminin begins to be expressed in S phase and persists through G2 to be degraded in G1 (**Figure 1, A and B**). Thus, interphase stages are labeled as G1 (Cdt1 only), early S (Cdt1 and Geminin), and late S/G2 (Geminin only). Furthermore, we tracked FUCCI in wild type with low nuclear blebbing and chromatin decompaction drugs known to induce increased nuclear blebbing (VPA and DZNep, **Figure 1C**). To determine if nuclear blebs enrich in a specific interphase stage, we compared the percentage of cells in each stage in the total population vs. blebbed nuclei. In wild type cells, the total percentage of cells was similar to the percentage of blebbed nuclei in each stage (**Figure 1D**), suggesting nuclear blebs do not prefer a specific interphase stage. Increased euchromatin via histone deacetylase inhibitor valproic acid (Gurvich *et al*., 2004) and decreased heterochromatin via histone methyltransferase inhibitor DZNep (Miranda *et al*., 2009) which increase nuclear blebbing, recapitulated similar distributions between total cells and cells with blebbed nuclei (**Figure 1, E and F**). Blebbed nuclei are present at an equal proportion to the population across the interphase cell cycle showing no preference in wild type or perturbed FUCCI cells.

**Figure 1.**
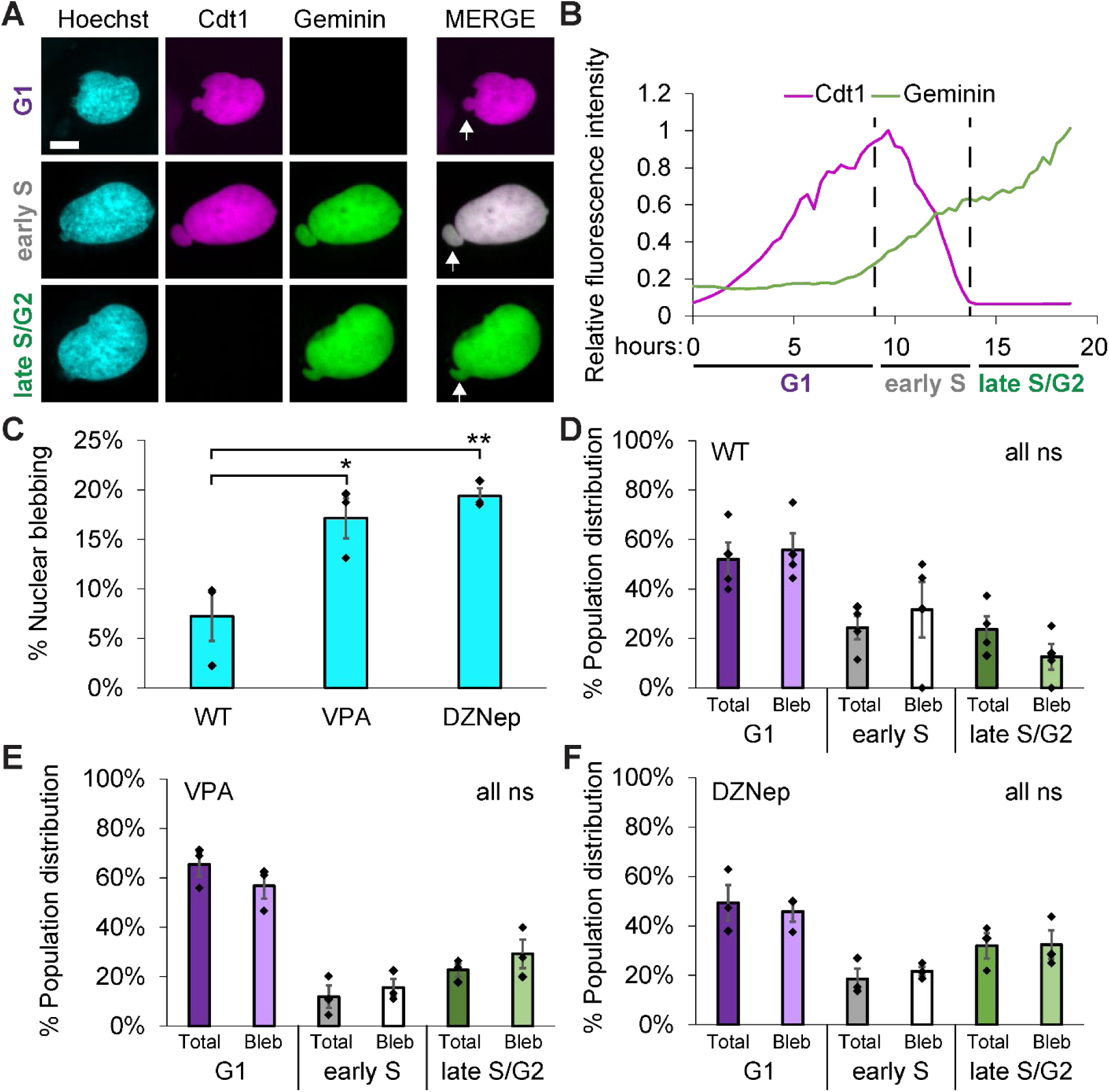
Static imaging of FUCCI cells reveals nuclear blebs are equally distributed throughout interphase stages. (A) Example images of HT 1080 FUCCI cells with Cdt1- and Geminin showing G1 (purple), early S (gray), and late S/ G2 (green). White arrow denotes nuclear bleb in merged image. (B) Example Graph of Cdt1 and Geminin intensity throughout the interphase part of the cell cycle for a single cell measured in hours. (C) Graph of percentage of cells displaying a nuclear bleb in wild type (WT) and chromatin decompaction via HDACi VPA or HMTi DZNep. Biological triplicates of n > 100 cells each. (D – F) Graphs of the percentage of cells in each interphase stage for the total population of cells (darker color) and for only cells presenting a nuclear bleb (lighter color) for (D) wild type, WT and for chromatin pertubations that increased nuclear blebbing (E) increased euchromatin via VPA and (F) decreased heterochromatin via DZNep. Biological replicates of 3 or 4 with a total of > 400 cells for population and > 20 cells with nuclear blebs. Unpaired two-tailed Student’s t-test p values reported as *<0.05, **<0.01, ***<0.001, or ns denotes no significance, p>0.05. Error bars represent standard error. Scale bar = 10 µm.

As an alternative way to measure interphase stage distributions in other cell types, we utilized BrdU incorporation to label nuclei in S phase. Again, we compared low nuclear blebbing in MEF wild type cells with higher nuclear blebbing upon chromatin perturbations (VPA and DZNep) or lamin perturbation (*Lmnb1-/-*, **Figure 2A**). BrdU via immunofluorescence imaging 30 minutes after BrdU addition labels nuclei actively replicating their DNA and thus in S phase (**Figure 2B**). The percentage of BrdU labeled nuclei in the total population and those with nuclear blebs were similar across all conditions. This agrees with Figure 1 FUCCI data, showing again that nuclear blebs are equally represented in S phase and non-S phase nuclei. Interestingly, between decreased heterochromatin perturbation (DZNep) and lamin B1 knockout (*lmnb1 -/-*) the percentage of total cells in S phase changed significantly, which aggress with reports that lamin B1 loss elongates S phase (Camps *et al*., 2014). Even though the whole population changes, the percentage of nuclei with blebs in S phase matches the population. This data suggests nuclear blebbing occurs equally across the interphase stages even when the distribution of cells undergoing DNA replication changes. HT1080 FUCCI cells labeled with BrdU showed that the percentage of total cells vs. percentage of blebbed nuclei in S phase was similar (**Figure 2D**). Taken together this data across two cell types and two methods reveals that nuclear blebs are presented equally throughout interphase stages.

**Figure 2.**
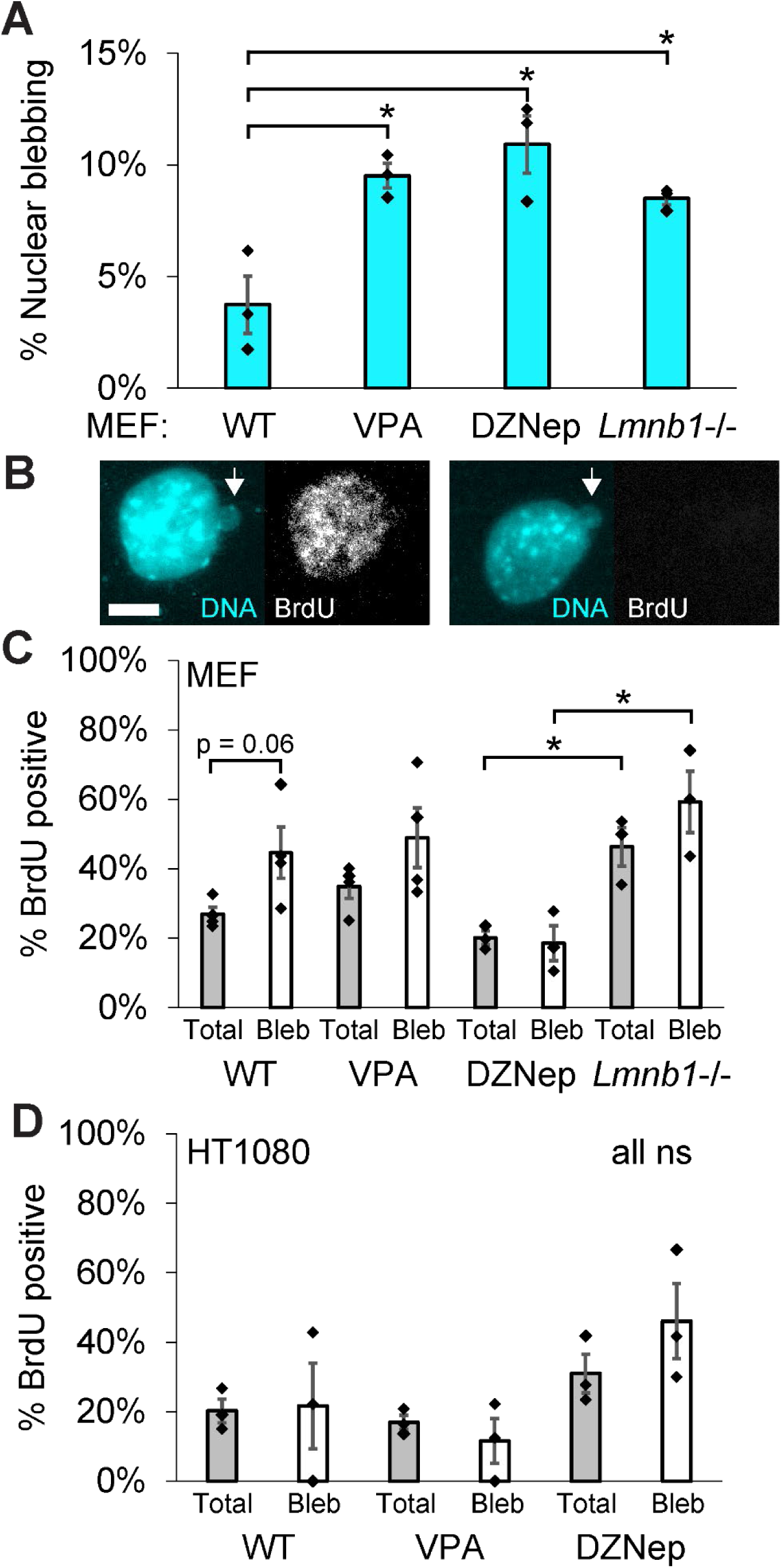
Nuclear blebs are present at population levels in replicating cells. (A) Graph of percentage of cells displaying nuclear blebbing in MEF wild type (WT) and perturbations VPA, DZNep, and loss of lamin B1(lmnb1-/-). Biological triplicates of n > 140 cells each. (B) Example images of a blebbed nucleus positive (left) and negative (right) for DNA analog BrdU that denotes cells undergoing replication. White arrow denotes nuclear bleb. (C) Graph of the percentage of MEF total cells (gray) and blebbed nuclei (white) that are positive for BrdU staining, and thus in S phase. (D) Graph of the percentage of HT 1080 total cells (gray) and blebbed nuclei (white) that are positive for BrdU staining, and thus in S phase. Biological replicates of 3 or 4 with each replicate containing n > 160 cells each and n > 7 blebs for WT and n > 17 blebs for every other condition. Unpaired two-tailed Student’s t-test p values reported as *<0.05, **<0.01, ***<0.001, or ns denotes no significance, p>0.05. Error bars represent standard error. Scale bar = 10 µm.

### Nuclear blebs preferentially form in G1 and persist in S and G2

Nuclear blebs form during interphase, persist, and are the site of repetitive nuclear envelope rupture (Stephens *et al*., 2018, 2019b). Using time lapse imaging to track nuclear shape, we aimed to determine when nuclear blebs form during the interphase stages of the cell cycle. FUCCI cells were imaged for 36 hours to ensure tracking cells from start to finish of interphase (**Figure 3A**). Time lapse imaging reveals that nuclear blebs form preferentially in G1 because at any given time 50% of cells are in G1 but 80% of blebs form in G1, a statistically significant increase and clear overrepresentation above population behavior for both wild type and VPA-based chromatin decompaction (**Figure 3A-C, Video 1**). Most nuclei that form a nuclear bleb rupture soon after (85% rupture 17/20 blebs MEF WT), as previously reported (Stephens *et al*., 2019b). These blebs persist into early S and late S/G2 reported by FUCCI fluorescence (**Figure 3A**) indicating that equal distribution of nuclear blebs across the cell cycle stems from the formation of a bleb in G1 and its persistence throughout S and G2 phases.

**Figure 3.**
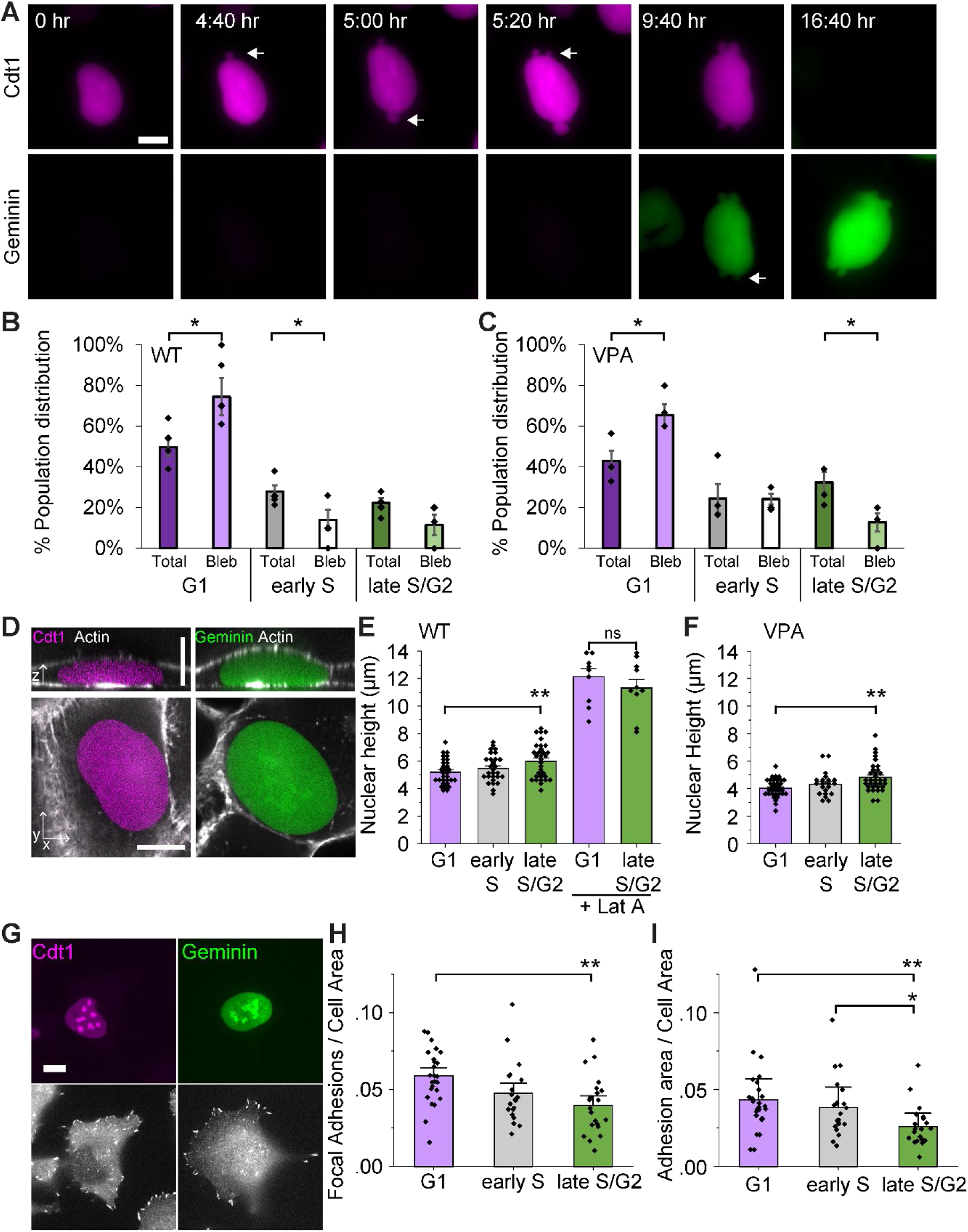
Time lapse imaging reveals that nuclear blebs form predominantly in G1 due to actin confinement and then persist throughout interphase. (A) Example images of a HT1080 FUCCI nucleus forming a blebs, denoted by arrows, in G1 and those blebs persisting into late S/G2. Time in hours (hr) throughout interphase denoted in white in the upper left of each image. (B, C) Graphs of the percentage population of total cells (darker) and cells in which a nuclear bleb formed (lighter) in G1, early S, or late S/G2 for both (B) wild type, WT and (C) chromatin decompaction via VPA. 4 and 3 biological replicates respectively where each replicate contains n > 180 cells and n > 8 nuclear blebs. MEF and RPE1 cells showed similar results for increased bleb formation relative to population, see **Supplemental** Figure 1. (D) Example images of nuclear height in G1 (purple) and late S/G2 (green) to measure actin confinement. (E, F) Graph of nuclear height for G1, early S, and late S/G2 in (E) wild type, WT without or with actin depolymerization drug latrunculin A (+ Lat A) for 1 hour and (F) VPA. Graphs are an average of n = 31,28, 32 respectivley for wild type (WT) G1, early S and late S/G2 without LatA and with LatA G1 and late S/G2 are both n= 10, and n = 34, 21, and 33 respectively for VPA-treated nuclei. RPE1 nuclei showed similar actin height changes from G1 to late S/G2, see **Supplemental** Figure 1. (G) Example images of G1 and late S/G2 nuclei with immunofluorescence of paxillin focal adhesions. Graphs of paxillin (H) focal adhesions per cell area and (I) adhesion area per cell area in wild type nuclei in G1, early S, and late S/G2, n = 24,20,21 respectively. Actin contraction was similar throughout interphase see **Supplemental** Figure 2. Unpaired two-tailed Student’s t-test p values reported as *<0.05, **<0.01, ***<0.001, or ns denotes no significance, p>0.05. Error bars represent standard error. Scale bar = 10 µm.

We time lapse imaged other cell lines to determine if over-representation of nuclear bleb formation in G1 could be recapitulated. Imaging MEF NLS-GFP nuclei normalized to time in the cell cycle recapitulates the formation of nuclear blebs occur predominantly in the first 50-60% (mostly G1) portion of the interphase cell cycle and then persists throughout interphase (**Supplemental Figure 1A**). Time lapse imaging of RPE1 FUCCI cells shows a similar trend as HT1080, with nuclear blebs forming at a higher frequency in G1 than the general cell population (**Supplemental Figure 1B**). Thus, dynamic imaging of nuclear bleb formation in multiple cell types consistently showed an increased frequency of forming a nuclear bleb in G1 that persists into other interphase cell cycle stages.

### G1 nuclei are under greater actin confinement than late S/G2

Actin confinement and actin contraction are two major factors that antagonize nuclear shape and cause nuclear blebbing (Le Berre *et al*., 2012; Hatch and Hetzer, 2016; Cho *et al*., 2019; Mistriotis *et al*., 2019; Nmezi *et al*., 2019; Xia *et al*., 2019; Pho *et al*., 2023). Thus, we assayed actin confinement by nuclear height and actin contraction by active actomyosin via phosphorylated myosin light chain 2 (pMLC2). Confocal imaging of live wild type HT 1080 FUCCI cells revealed that nuclear height in G1 nuclei 5.2 ± 0.2 µm was significantly decreased (actin confinement increased) relative to taller late S/G2 nuclei 6.0 ± 0.2 µm for both wild type and VPA-treated **(Figure 3, D and E**). To directly test whether actin confinement determines nuclear height, cells were treated with actin depolymerization drug latrunculin A for 1 hour, which drastically increased nuclear height and abolished the differences between G1 and late S/G2 nuclei (**Figure 3E**, +Lat A). HT1080 FUCCI VPA-treated cells show a decreased G1 nuclear height relative to wild type from 5.2 to 4.0 ± 0.1 µm, which agrees with previous reports of decreased nuclear height (increased actin confinement) and is consistent with reported decreased chromatin rigidity (Berg *et al*., 2023; Pho *et al*., 2023). These more confined VPA-treated cells recapitulated the increased nuclear height from G1 nuclei at 4.0 ± 0.1 µm to late S/G2 nuclei at 4.8 ± 0.2 µm (p< 0.001, **Figure 3F**), denoting decreased actin confinement. RPE1 FUCCI cells recapitulated shorter G1 nuclei (more actin confinement) and taller late S/G2 nuclei (less actin confinement, **Supplemental Figure 1C**). This data agrees with reports of increased actin confinement in G1 from others (Aureille *et al*., 2019). Measurements of actin contraction via pMLC2 reveal no significant change in active actomyosin between the different stages of interphase (**Supplemental Figure 2**). Thus, G1 nuclei are under increased actin confinement compared to late S/G2 nuclei, while actin contraction remains similar across the interphase stages.

Actin confinement relies on the cell attaching by focal adhesions to the substrate (Kim *et al*., 2012) and these focal adhesions are removed in G2 in preparation for cell rounding in mitosis (Jones *et al*., 2018). To quantify the number of focal adhesions in each interphase stage, we fixed FUCCI cells and performed immunofluorescence for the focal adhesion protein paxillin (**Figure 3G**). G1 nuclei measured a statistically significant increase in both focal adhesion density and area relative to cell size compared to late S/G2 nuclei (**Figure 3, H and I**). Thus, G1 have more focal adhesions providing a plausible mechanism for increased actin confinement leading to increased nuclear bleb formation.

### Artificial compression shows that nuclei in G2 are more fragile than in G1

One hypothesis is that G1 nuclei, which natively are more likely to form a bleb, will show more nuclear ruptures under increased artificial confinement. An alternative hypothesis is that G2 nuclei will rupture more because they are weaker but usually under less confinement due to alleviation of focal adhesions. To test these hypotheses directly, we artificially modulated confinement using a compression device using a coverslip with micropillars of defined heights (Le Berre *et al*., 2012). This experiment provides the ability to compress nuclei of cells in different interphase stages to the same height.

FUCCI cells and artificial confinement provide the ability to visualize how neighboring asynchronous cells from different interphase stages respond to specific confinement heights. Wild type measurements reveal that G1 nuclei are 5.2 ± 0.2 µm in height (**Figure 3E**). Thus, we placed the nuclei under increased artificial confinement to 4 µm (**Figure 4A**). Under artificial confinement of 4 µm G1 nuclei displayed nuclear rupture in 17 ± 5% of cells (**Figure 4B** **and Video 2**). However, late S/G2 nuclei displayed a significant increase in nuclear rupture under the same 4 µm artificial confinement to 49 ± 8%. This difference in rupture percentages was not due to changes in percent size increase in x and y dimensions upon confinement (p > 0.05, **Figure 4C**). To show that confinement does determine nuclear rupture, we then placed nuclei under increased artificial confinement to 3 µm resulting in increased nuclear rupture relative to 4 µm confinement for both G1 and late S/G2 nuclei (**Figure 4B**). Upon this drastic increase in artificial confinement G1 and late S/G2 nuclei showed a statistically similar 85-96% nuclear rupture. Furthermore, upon applying 3 µm artificial confinement, late S/G2 nuclei ruptured before neighboring G1 nuclei (**Figure 4D**). Thus, late S/G2 nuclei are weaker than G1 nuclei under similar confinement.

**Figure 4.**
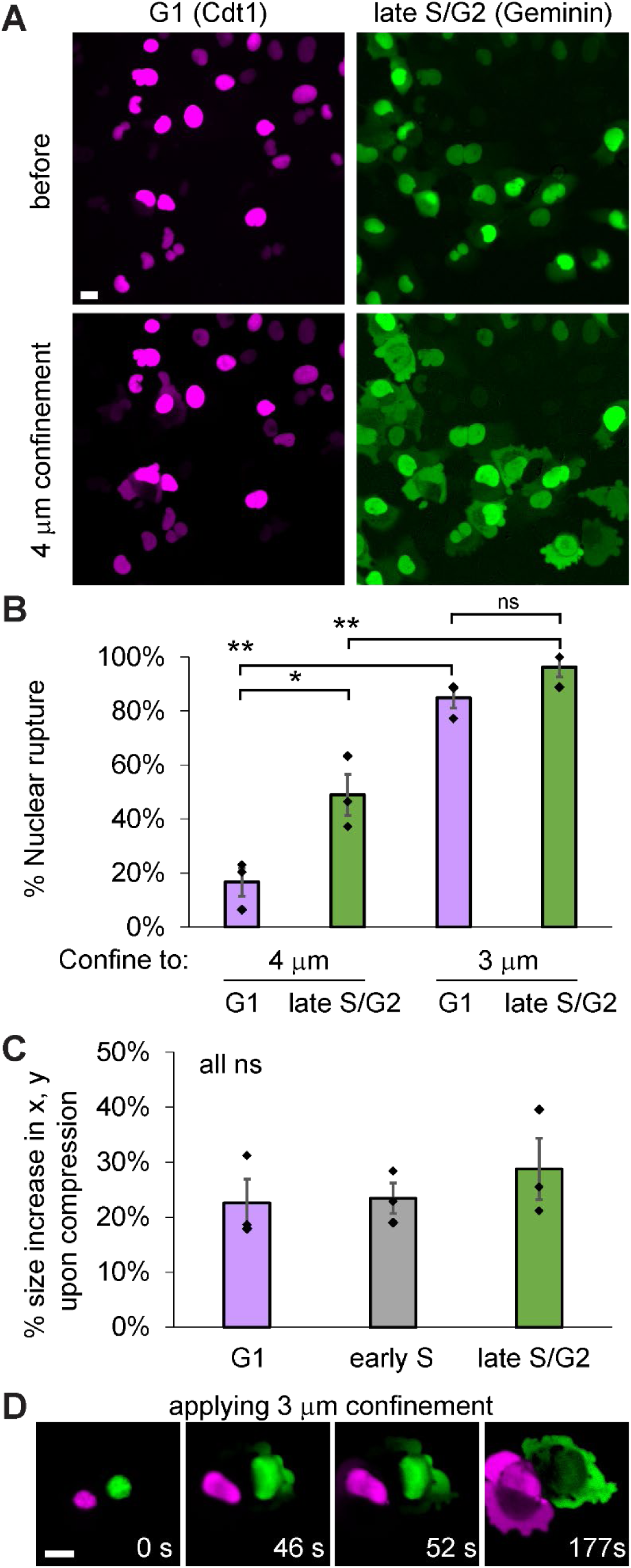
Late S/G2 nuclei rupture more frequently than G1 nuclei when placed under artificial confinement. (A) Example images of a wild type HT 1080 FUCCI labeled nuclei via Cdt1 (only = G1) and Geminin (only = late S/G2) before and after 4 µm artificial confinement. (B) Graph of the percentage of G1 and late S/G2 nuclei that rupture upon artificial confinement to 4 µm and 3 µm. Triplicates each with n > 89 cells for 4 µm and n > 30 cells for 3 µm artificial confinement. (C) Example image of cells progressively undergoing artificial confinement to 3 µm over time shows that late S/G2 nucleus ruptures before the neighboring G1 nucleus. Time denoted in white as seconds, s. (D) Graph of the percentage increase in nuclear size measured in x, y upon 4 µm artificial confinement in wild type HT 1080 cells in G1, early S, and late S/G2. Unpaired two-tailed Student’s t-test p values reported as *<0.05, **<0.01, ***<0.001, or ns denotes no significance, p>0.05. Error bars represent standard error. Scale bar = 10 µm.

### Micromanipulation force measurements confirm G2 nuclei are weaker than G1

Nuclear blebbing and rupture stems from loss of nuclear stiffness from chromatin decompaction and/or loss of lamins (Kalukula *et al*., 2022). To aid nucleus isolation from the cytoskeleton for dual micropipette micromanipulation force measurements, we use MEF vimentin null nuclei (Stephens *et al*., 2017; Currey *et al*., 2022). By isolating a single nucleus and puling on it via micromanipulation while measuring forces, we can separate the nuclear mechanical contributions of chromatin and lamins. During micromanipulation the pull pipette extends the nucleus while deflection of the force pipette multiplied by its pre-measured bending constant provides a measure of force. Chromatin rigidity determines the short extension regime (0-3 µm extension) while lamins determine strain stiffening that occurs at higher extensions > 3 µm (Banigan *et al*., 2017; Stephens *et al*., 2017; Currey *et al*., 2022). This force vs. extension relation can be graphed with slope providing a nuclear spring constant nN/µm (**Figure 5**).

**Figure 5.**
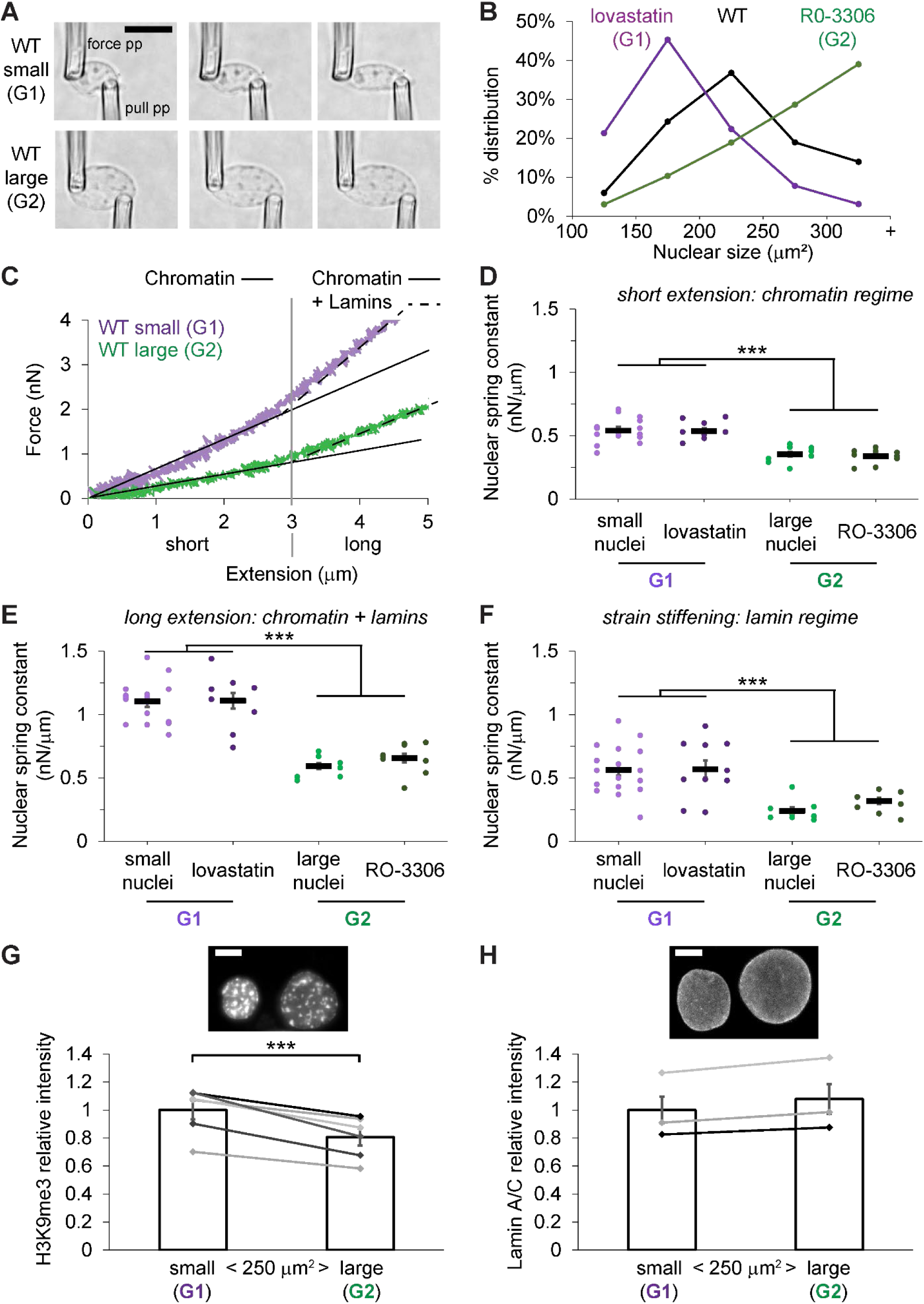
Micromanipulation nuclear force measurements confirm that G2 nuclei are mechanically weaker than G1 nuclei. (A) Example images of a small G1 and large G2 nuclei during micromanipulation force extension measurements. The pull pipette extends the nucleus as deflection of the force pipette multiplied by its pre-measured bending constant provides a measure of force. Scale bar = 10 µm. (B) Graph of percentage distribution of total population by nuclear size for G1 stall via lovastatin (purple, n = 188), wild type (WT, black, n = 632), and G2 stall via R0-3306 (green, n = 186). (C) Example graphs of force in nanonewtons vs. extension in micrometers measurement shown for small and large MEF vimentin null nuclei representing G1 and G2 respectively. The slope of the line provides a nuclear spring constant (nN/ µm) averaged for each nucleus over three force vs. extension pulls. The short extension (< 3 µm) measures chromatin’s mechanical contribution (solid line), where long extension (> 3 µm) shows strain stiffening due to lamins (dashed line). (D-F) Graphs of nuclear spring constants for (D) Chromatin-based short extension, (E) long extension which accounts for both chromatin and lamin contribution, and (F) lamin-based strain stiffening at longer extensions. Nuclear spring constants are reported for wild type small nuclei, G1 stall via lovastatin, wild type large nuclei, and G2 stall via R0-3306 (n = 17, 11,8, 11). (G,H) Example images and graphs of paired small and large nuclei relative immunofluorescence intensity for (G) constitutive heterochromatin marker H3K9me3 and (H) lamin A/C (N = 6 H3K9me3 and 3 lamin A/C biological replicates where each contained > 20 nuclei for each condition). Unpaired (D-F) or paired (G and H) two-tailed Student’s t-test p values reported as *<0.05, **<0.01, ***<0.001, or ns denotes no significance, p>0.05. Error bars represent standard error. Scale bar = 10 µm.

To determine which nuclei were in G1 and G2, we measured nuclear size which relatively doubles. We compared nuclear size distributions upon use of drug inhibitors to stall the cell cycle (Iida *et al*., 2022), which shifted to smaller nuclei for G1 stall via lovastatin and larger nuclei for G2 stall via RO-3306 compared to wild type (**Figure 5B**). Using this data in asynchronous cell culture we selected smaller nuclei 100-200 µm^2^ size to denote G1 and larger nuclei >300 µm^2^ size to denote G2. Relative to smaller nuclei, larger nuclei were significantly weaker measuring a decrease in the chromatin dominated short regime nuclear spring constant from 0.54 ± 0.03 nN/µm in small nuclei to 0.35 ± 0.02 nN/µm in large nuclei (**Figure 5, C and D**). Interestingly, long extension nuclear spring constant which is composed of both chromatin and lamin contributions also significantly decreased in larger nuclei from 1.10 ± 0.04 to 0.60 ± 0.03 nN/µm, **Figure 5E**). This decrease could be due to the large contribution of chromatin-based rigidity. To measure lamin-based strain stiffening, we subtracted the short from the long extension nuclear spring constant. Lamin-based strain stiffening also decreased from small to large nuclei (0.56 ± 0.03 to 0.24 ± 0.03 nN/µm, **Figure 5F**). Thus, both chromatin- and lamin-based contributions to nuclear rigidity decreased from stronger smaller nuclei representing G1 to weaker larger nuclei representing G2, which confirms the results from the artificial confinement experiment where G2 nuclei ruptured at a great frequency than G1 under the same confinement.

Next, we used cell cycle stalling drugs to synchronize cells in G1 via lovastatin and G2 via R0-3366. We performed a similar set of micromanipulation force measurements which returned near identical values for the asynchronous small and larger nuclei used as proxies for G1 and G2. Nuclei from lovastatin treated cells to cause a G1 stall measured a statistically similar stronger short, long, and strain stiffening extensions as compared small nuclei. Nuclei from R0-3366 treated cells to stall in G2 measured a statistically similar weaker short, long, and strain stiffening extensions similar values for large nuclei. Thus, relative to lovastatin G1 stalled nuclei, RO-3366 G2 stalled nuclei were significantly weaker in the chromatin-based short extension, chromatin and lamin long extension, and lamin-based strain stiffening nuclear spring constants (**Figure 5D-F**). Taken together, we provide novel evidence that G2 nuclei are weaker than G1 nuclei due to weakening of both major nuclear stiffness components, chromatin and lamins.

To determine the source of changes in both the chromatin and lamin nuclear spring constants, we measured chromatin histone modifications and lamin levels via immunofluorescence. The constitutive heterochromatin marker H3K9me3 decreased from G1 to G2 nuclei (**Figure 5H**) while lamin A/C levels remined similar (**Figure 5H**). We recapitulated these findings in HT1080 FUCCI human cells where constitutive heterochromatin decreases lamin A/C remains the same from G1 to late S/G2 (**Supplemental Figure 3**). Thus, the decrease from G1 to G2 in chromatin-based nuclear stiffness could be due to decreased H3K9me3 but not the decreased in lamin-based nuclear stiffness remains unknown.

### Loss of H3K9me3 can account for decreased chromatin and lamin nuclear stiffness from G1 to G2

Chromatin is known to connect with lamins in cells (Harr *et al*., 2015; van Schaik *et al*., 2020) and biophysical modeling suggests these connections are mechanical linkers necessary for the two regime behavior of the nucleus via physics modeling (Banigan *et al*., 2017; Strom *et al*., 2021). Constitutive heterochromatin marker H3K9me3 binds to lamins, has peripheral localization, and also shows internal localization to chromocenters which are dense chromatin foci ((Manning *et al*., 2025), **Figure 6A**). Using spinning disk confocal imaging, we find that relative to smaller nuclei (G1), larger nuclei (G2) have both decreased peripheral and whole nucleus H3K9me3 levels (**Figure 6A-C**). The loss of H3K9me3 specifically from G1 to G2 could result in loss of chromatin-lamin linkers previously modeled to affect the lamin strain stiffeneing response (Strom *et al*., 2021). Analysis of chromocenters shows that the intensity and size of chromocenters is similar from small to large nuclei (**Figure 6D**). Thus, loss of periphery and whole nucleus H3K9me3 could account for decreased nuclear stiffness from both the chromatin and lamin regime from G1 to G2.

**Figure 6.**
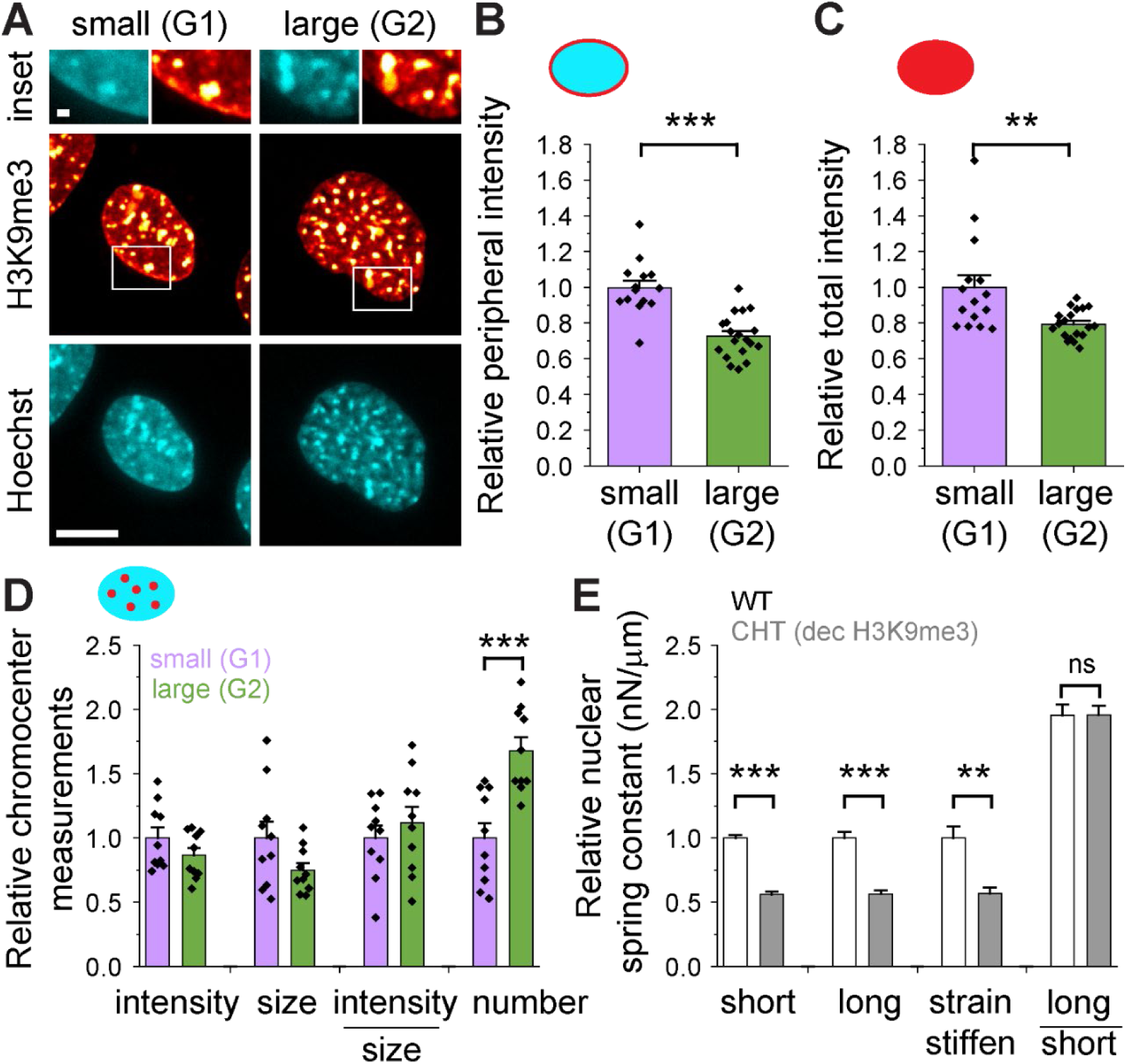
Loss of H3K9me3 occurs in the interphase cell cycle and causes a decrease in both chromatin- and lamin-based nuclear mechanics. (A) Example confocal images of a nucleus stained for DNA via Hoechst (cyan) and constitutive heterochromatin H3K9me3 (red). Inset highlights the differences in peripheral H3K9me3. Graphs of H3K9me3 relative (B) peripheral and (C) whole nucleus intensity for smaller (100-200 µm^2^) and larger (>300 µm^2^) nuclei representing respectively G1 and G2 nuclei (n = 15 and 19). Similar decreases in whole nucleus and periphery H3K9me3 are also seen in HT1080 cells in **Supplemental** Figure 3C. (D) Graph of H3K9me3 relative chromocenter intensity, size, intensity/size, and number for smaller (100-200 µm^2^) and larger (>300 µm^2^) nuclei representing respectively G1 and G2 nuclei (n = 10 and 10). (E) Graph of relative nuclear spring constant single nucleus micromanipulation force measurement for wild type and SUV39H1 inhibitor Chaetocin to decrease H3K9me3. These measurements were made using previously published data (Manning *et al*., 2025). Relative comparisons for each nuclear stiffness regime short (chromatin-based), long (chromatin + lamin A regime), strain stiffening (lamin-based, long – short), and ratio of long vs. short (n = 13 and 10 respectively). Unpaired two-tailed Student’s t-test p values reported as *<0.05, **<0.01, ***<0.001, or ns denotes no significance, p>0.05. Error bars represent standard error. Scale bar = 10 µm.

H3K9me3 levels can be decreased via SUV39H1 methyltransferase inhibitor chaetocin (Greiner *et al*., 2005). We recently showed that this loss of H3K9me3 decreases of both chromatin and lamin-based nuclear stiffness in smaller G1-like nuclei (Manning *et al*., 2025). To show this effect, we reanalyzed the data in the previous study as relative change in nuclear spring constant. This analysis indicates that loss of H3K9me3 via Chaetocin decreases the short extension, long extension, and strain stiffening (long – short) regimes (**Figure 6E**) which recapitulated nuclear spring constant changes from G1 to G2 (**Figure 5**). This data is consistent with mechanical modeling of loss of chromatin to lamin linkages, which predicts deceases in both the chromatin short extension regimes and the lamin long extension strain stiffening (Strom *et al*., 2021). Taken together the data show that the measured loss of H3K9me3 from G1 to G2 nuclei is sufficient to alter both the chromatin and lamin-based nuclear stiffness regimes.

## Discussion

### G1 nuclei are both stronger and under greater actin antagonism

Our data reveal that nuclear blebs form in G1 at an over-represented rate relative to the total population of cells that leads to nuclear ruptures which causes cellular dysfunction. Moreover, interphase nuclear and actin mechanics interaction results in the force balance necessary to maintain nuclear homeostasis. Thus, changes in either nuclear mechanics and/or actin or external confinement can result in increased nuclear bleb formation in G1/G0 nuclei. In theory, the ability for G1 nucleus to form a nuclear bleb might start the cell and nucleus down a path of dysfunction that contributes to a worsening disease state.

Nuclear confinement is sufficient to overcome a stronger G1 nucleus to cause nuclear blebbing and ruptures. Nuclear confinement via actin can overcome even the strongest nucleus to cause nuclear blebs to form, as evidenced by the disproportionate increase in nuclear bleb formation in G1 nuclei. From our previous work, the strength of actin confinement can be seen in chromatin decompacted nuclei via VPA, in which the actin compresses the nuclear height significantly by 0.5 um, or ∼10% of overall nuclear height, due to the weaken of the chromatin-based nuclear stiffness (Berg *et al*., 2023; Pho *et al*., 2023). The importance of nuclear confinement is further confirmed by artificial compression (**Figure 4**). A change from 4 µm artificial confinement to 3 µm confinement resulted in a significant increase in nuclear ruptures to near 100% (84-96%) of all nuclei to deform and rupture, independent of nuclear interphase stage and strength. Thus, our data suggests that even a strong nucleus will deform and rupture under high levels of actin confinement.

### S phase DNA replication does not affect nuclear blebbing but might impact dysfunction from nuclear blebbing

The action of DNA replication has no significant impact on nuclear blebbing. Our data clearly show that nuclear blebs are present at and form at population distribution levels, or below, showing no support for DNA replication having a role in nuclear bleb formation. While DNA replication has no measurable impact on nuclear blebbing, it has been reported to impact dysfunction associated with nuclear blebbing and rupture via DNA damage. The literature remains contentious for the importance of nuclear rupture to cause increased DNA damage. Many studies have shown that deformation of the nucleus without rupture is sufficient to induce increased DNA damage (Denais *et al*., 2016; Raab *et al*., 2016; Xia *et al*., 2018). However, deformation only or rupture increased DNA damage have both been reported to be reliant on DNA replication for these increased levels of DNA damage(Irianto *et al*., 2017; Shah *et al*., 2021; Chu *et al*., 2025).

While DNA replication and its associated S phase have no apparent role, the result of going through this process leads to G2 and its effects on both nuclear and actin mechanics. One hypothesis is that cancer cells that are constantly going through the cell cycle might be more susceptible to nuclear blebbing and rupture due to passing through the interphase stages of DNA replication (S phase) and onto G2. Thus, S phase does not significantly alter nuclear blebbing but it does lead to a phase that does impact nuclear and actin force balance determining nuclear integrity and thus function.

### G2 nuclei are mechanically weaker but under less actin confinement

We find overall that G2 nuclei are weaker and thus lack the ability to maintain nuclear shape and integrity in the face of actin and/or external confinement. This finding has strong implications in human diseases where cells might be cycling and thus be in S and G2 more often than non-cycling cells in a G1/G0 state. Our findings reveal that G2 nuclei are more fragile under artificial confinement (**Figure 4**) which is supported by decreased nuclear stiffness via dual pipette micromanipulation (**Figure 5**). In agreement, other publications have also reported similar findings. In glioblastoma human cell lines, U251MG blebs and ruptures more than U87MG. U251MG has no p21 and is thus cycling more than G1 U87MG (Kamikawa *et al*., 2023). This paper concludes that G1 cells are more resistant to nuclear envelope stress than S/G2.

Nuclear stiffness decreases for both the chromatin and lamin regimes. Our direct data shows this change in chromatin stiffness is partially due to loss of constitutive heterochromatin marker H3K9me3. However, the data is less clear about why lamin-based strain stiffening is decreased because there is no significant loss of lamin A/C. Chromatin-lamin interactions are the other sub-component of nuclear force response. Physics modeling of nuclear mechanics shows that loss of chromatin-lamin interactions would result in changing both chromatin and lamin-based nuclear stiffness (Strom *et al*., 2021). H3K9me3 interacts with the lamina through PRR14 tethering and HP1a. H3K9me3 nuclear periphery enrichment is significantly lost from G1 to G2, suggesting a mechanism for strain stiffening-based mechanical changes. Upon depletion of H3K9me3 via a SUV39H1 inhibtor Chaetocin, which mimics loss whole nucleus as well as periphery enrichment of H3K9me3, nuclear stiffness is lost in both the chromatin short regime and lamin-based long extension strain stiffening (**Figure 6**). Thus, loss of heterochromatin which acts as a chromatin-lamin linker is sufficient to recapitulate changes in nuclear stiffness from G1 to G2.

While we provide a plausible mechanism for cell cycle-based decreased nuclear stiffness other factors might also contribute to the loss of chromatin-lamin linkers. DamID data reveals that lamin associated domains have cell cycle specific behaviors (van Schaik *et al*., 2020). Telomere proximal chromatin is associated with the lamina in G1 but detaches over the cell cycle while centromere proximal chromatin accumulates over the cell cycle. It was hypothesized that changes in telomere or end chromosome association detachment was due to changes in LAP2α (Dechat *et al*., 2004). Overall, similar to loss of focal adhesions in lieu of cell rounding in mitosis, loss of chromatin to lamin connections makes sense to both condense the chromatin into chromosomes and prepared for nuclear envelope breakdown at the onset of mitosis.

One of the nuclear stiffness sub-components is chromatin-chromatin linkages shown to be essential to nuclear mechanical response in simulations (Banigan *et al*., 2017; Strom *et al*., 2021). Our lab and others have shown that local chromatin linker HP1α is a key mechanical component determining nuclear shape (Strom *et al*., 2021; Williams *et al*., 2024). HP1α is reported to be phosporylated during the cell cycle to localize to kinetochore (Chakraborty *et al*., 2014). This might decrease HP1α across the genome leading weaking the chromatin gel polymer interior. We have also shown that long distance chromatin interactions via HiC provide chromatin mechanical stiffness to the nucleus (Belaghzal *et al*., 2021). To this point, cell-cycle dynamics of chromosomal organization at single-cell resolution shows that chromatin HiC contacts in G1 are mostly long range interactions at 50 Mb which dramatically shifts in G2 to shorter range interactions at 250 Kb (Nagano *et al*., 2016). Thus, it is possible that moving from long range to shorter range interactions might also contribute to decreased chromatin-based nuclear mechanics. Overall, it remains unclear how or why a duplicated genome would scale chromatin-chromatin interactions for a transient period of time before mitosis.

Similar but differently, actin focal adhesions are removed in lieu of cell rounding in mitosis. Decreased actin confinement in G2 nuclei results in less or at population levels nuclear bleb formation even though the nucleus is weaker in G2 (**Figure 3A-F**). Decreased actin confinement from G1 to G2 have been confirmed by others via Hela, MEF, and MRC-6 cells (Aureille *et al*.,

2019). This is further supported by decreased traction forces during cell cycle progression (Vianay *et al*., 2018). The mechanism underlying this phenomena is loss of focal adhesions from G1 to G2 (**Figure 3G-I**), which is supported by other publications measuring the same outcome (Jones *et al*., 2018). Previous work has established that focal adhesions control the actin cap confinement of the nucleus (Kim *et al*., 2012). For both actin and nucleus mechanics change during the cell cycle in a functional manner to move the cell into mitosis, but the these changes has real consequences in the integrity of the nucleus and its functions.

## Materials and Methods

### Cell culture

MEF WT NLS-GFP (*Lmnb1*-/- NLS GFP, *V-/-*), and HT1080 FUCCI were cultured in DMEM (Corning) containing 10% fetal bovine serum (FBS, HyClone) and 1% penicillin/streptomycin (Corning). RPE-1 FUCCI were cultured in DMEM/F-12 50/50 (Corning) complete with 10% FBS and 1% penicillin/streptomycin. The cells were incubated at 37 °C and 5% CO2 with humidity and passaged every 2-3 days for no more than 30 generations. MEF cells were obtained from the Goldman Lab (Department of Cell and Molecular Biology, Northwestern University Feinberg School of Medicine, Chicago, IL USA), and HT1080 and RPE-1 FUCCI cells were obtained from Orth Lab (Department of Molecular and Cellular and Developmental Biology, University of Colorado Boulder, Boulder CO USA).

### Drug treatments

Cells were treated with 4mM VPA (1069-66-5, Sigma) from a 20 mM dilution in complete DMEM. Cells were treated with 2.5 µM 3-deazaneplanocin (DZNep, 120964-45-6, Cal Biochem) from a 25 mM stock solution in cell culture grade water. Cells were treated with 10 µM RO-3306 (872573-93-8, Sigma) prepared from a 10 mM concentration in DMSO. Cells were treated with 10-25 µM lovostatin (H52792.06, Thermofisher) prepared from a 50 mM stock solution in DMSO. DZNep, RO-3306 and lovostatin stock solutions did not exceed more than two freeze thaw cycles. Cells were imaged after 16-24 hours of treatment with VPA and DZNep, 16 hours with RO-3306 and 24 hours with lovostatin.

### Immunofluorescence

Cells were plated in 8 well glass chambers (Cellvis) and treated as above. For BrdU experiments, after reaching 70% confluence, cells were treated with 0.03 mg/mL BrdU (B23151, Invitrogen) prepared from a 98 mM stock concentration in DMSO. Cells were treated and incubated with BrdU for 30 minutes prior to fixation in cold ethanol for 5 minutes. The cells were then washed three times with PBS (Corning) for 5 minutes, followed by denaturation with 1.5 M HCl (VWR) for 30 minutes. For all other immunofluorescence experiments, cells were fixed in a solution of 4% paraformaldehyde in PBS for 15 minutes and washed three times in PBS for 5 minutes. Cells were then washed two times with PBS followed by permeabilization with 0.1% Triton X-100 (US Biological) in PBS for 15 minutes at room temperature, followed by a wash with 0.06% Tween 20 (US Biological) in PBS for 5 minutes. Cells were washed three times with PBS for five minutes each, followed by blocking for one hour at room temperature in 2% bovine serum albumin (BSA) (9048-46-8, Fisher) in PBS. Primary antibodies were diluted in 2% BSA and incubated for 12 hours at 4° C. Primary antibodies used were: BrdU Mouse Mab at 1:1000 (59992s, Cell Signaling), pMLC2 rabbit Ab 1:100 (3672, Cell Signaling Technologies), Mouse Anti-Paxillin at 1:1000 (610052, BD Transduction Laboratories), H2K9ac 1:400 (9649, Cell Signaling Technologies), Lamin A/C 1:10,000 (4777, Cell Signaling Technologies), H3k9me^2-3^ 1:400 (5327, Cell Signaling Technologies), and Lamin B1 (1:1000, ab16048 Abcam). Cells were washed three times with PBS. The secondary antibodies used were Alexa Fluor 555 anti-mouse IgG (4409S, Cell Signaling Technology), Alexa Fluor 647 anti-mouse IgG (4410S, Cell Signaling Technologies), Alexa Fluor 647 anti-Rabbit IgG (4414, Cell Signaling Technologies). Cells were treated with secondary antibodies 1:1000 in blocking solution on the shaker for one hour at room temperature in the dark. Cells were then washed 3 times in PBS before staining with a 1 µg/mL dilution of Hoechst 33342 (Life Technologies) in PBS for 15 minutes followed by three washes in PBS. Cells were mounted using ProLong Gold antifade (Life Technologies) and cured for 12 hours at room temperature in the dark. To label RNA, EU incorporation was detected using the Click-iT RNA Alexa Fluor 594 Imaging Kit (C10330). As previously described (Berg et al. 2023). 1 mM EU was added for 1 hour prior to fixation. Following permeabilization, 500 µl of the Click-iT reaction mixture was added and incubated for 30 minutes in the dark, followed by a rinse in the Click-iT reaction rinse buffer.

### Imaging

Nikon Elements software was used to acquire images on a Nikon instruments Ti2-E inverted widefield microscope, Orca Fusion Gen III camera, Lumencor Celesta light engine, TMC CleanBench air table, with 40x air objective (N.A. 0.75, W.D. 0.66, MRH0041), Plan Apochromat Lambda 100x oil immersion objective lens (N.A. 1.45, W.D. 0.13 mm, MRD71970). For height measurements and imaging of paxillin, a Crest V3 Spinning Disk Confocal was used. Imaging of cell confinement was acquired on a Nikon Ti2 inverted widefield microscope using a Plan Apochromat Lambda 20x air objective N.A. 0.75 (Nikon) with a Prime BSI-Express camera (Photometrics). Nikon Elements software was used for analysis, exported to Excel or Origin, and statistical significance was determined using unpaired two-tailed Student’s T-test (*p<0.05, **p<0.01, ***p<0.001). Not added: MM imaging

### Immunofluorescence imaging and analysis

Regions of interest were selected to obtain mean intensities of every nuclei. The signal to noise ratio was calculated using mean intensity signal from the ROIs and background taken from every image using a 15×15 pixel square ROI region with no cells. A signal to noise ratio greater than 1.5 was the threshold used to identify BrdU positive cells. The cells containing nuclear blebs were compared to the total population by the percent BrdU positive nuclei. To analyze RNA labeling, Z stacks were compiled into a maximum intensity projection and background was subtracted from a 15×15 pixel square region containing no cells. Regions of interest were drawn around single nuclei and the mean intensities of RNA labeled EU was collected from 3 fields of view. Add euchromatin, heterochromatin, pMLC Add image details bit, power, exposure time

Focal adhesion complexes were measured using paxillin immunofluorescence of HT1080 FUCCI cells by confocal imaging with 100x oil immersion lens in 12 bit sensitive between 200-300ms and 40-50% power, with Focal adhesion complexes were quantified by number of focal adhesions per cell area and sum adhesion area per cell area. Background was subtracted from a 3×3 µm square ROI.

### Live cell imaging and analysis

Cells were grown on 4 well chamber glass dishes (Cellvis) and treated as above. Cells were treated with 1 µg/mL Hoechst 33342 (Life Technologies) for 15 minutes prior to live cell imaging. Exposure times for Hoechst (DAPI), cdt1-RFP (TRITC), geminin-GFP (FITC) were between 50-200 ms in 12 bit. Blebbing was calculated by counting the total number of blebs/total nuclei for every replicate. Time lapses were acquired using a humidity chamber complete with Okolab heat and 5% CO2. Images were saved using NIS Elements AR and data was collected in Excel. Nuclear rupture was determined as outlined previously quantitatively defined a nuclear rupture as a > 25% increase in the cell/nucleus NLS-GFP intensity ratio (Pho *et al*., 2023).

### Analysis of Fluorescent Ubiquitin Cell Cycle Indicator (FUCCI)

To measure the interphase stage using FUCCI, regions of interest were drawn around nuclei and mean intensities of Cdt1-RFP and Geminin-GFP were collected using NIS Elements Analysis, and exported to Excel. Background from a 15×15 pixel area in a region where there are no cells was collected from every frame to generate signal to noise ratios of cdt1-RFP and geminin-GFP. A signal to noise greater than 1.5 was considered signal, where interphase stages were defined as G1 (cdt1-RFP positive, geminin-GFP negative), early S (cdt1-RFP positive, geminin-GFP negative), and late S/G2 (cdt1-RFP negative, geminin-GFP positive).

### Live cell imaging of nuclear height and analysis

Cells were grown in 4 well chamber glass dishes (Cellvis) and were untreated or treated 4 mM VPA the day after plating (1069-66-5, Sigma). Cells were stained with 1 µg/mL Hoechst 33342 (Life Technologies) for 15 minutes prior to imaging. Height images were taken on Nikon Eclipse Ti2 microscope with a 100x oil objective lens with Crest V3 Spinning Disk Confocal. Images were taken as 77 0.2 µm steps in 12 bit sensitive with exposure times for Hoechst (DAPI), Cdt1-RFP (TRITC), and geminin-GFP (FITC) were between 50-100 ms. As described previously (Pho et al 2023) 2 intensity line scans were drawn approximately equal in distance from the nucleus center through the Z plane of each nuclei, values were exported to Excel, and the average of the full-width half-max of the two line scans was calculated.

### Live Cell imaging of cell confinement and analysis

To confine the cells, we used a one-well dynamic cell confiner with a suction cup, and confiner slides of 3 or 4 microns (4D cell). We hydrated the coverslip and suction cup in complete DMEM one hour before use. Confinement was gradually applied by decreasing the pressure from -2 kPa to −10 kPa. Images were taken in FITC and TRITC in 16-bit between 4-10% power and 100-200 ms. Loss of compartmentalization of the nucleus was measured as a 20% loss of signal to noise of an ROI drawn before and after confinement.

### Micromanipulation Force Measurements

As previously described (Stephens *et al*., 2017; Currey *et al*., 2022), MEF Vimentin null (MEF *V-/-*) were grown in a micromanipulation well. Nuclei were isolated from living cells using a spray micropipette containing Triton X-100 (0.05%) in PBS. A pull micropipette was used to grab the nucleus, while the opposite end of the nucleus was grabbed by a precalibrated force micropipette and suspended in preparation for force-extension measurements. The pull pipette was moved 50 nm/s to extend the nucleus 3 or 6 µm. Nucleus extension was measured by tracking the pull micropipette (µm), and force (nN) was measured by deflection of the force micropipette multiplied by the bending modulus (1.2-2 nN/ µm). The slope of the force versus extension plot provides the spring constant (nN/µm) for the short chromatin-dominated regime (<3 µm) and long-extension lamin A-dominated strain-stiffening regime (>3 µm). The long-regime spring constant minus the short-regime spring constant provides the measure of lamin A-based strain stiffening. Size thresholds separating interphase stages are reported in Figure 5.

## Supplemental Figures

Supplemental Figure 1 provides recapitulation of increased nuclear bleb formation early in the interphase stages (G1) for both MEF and RPE-1 cells. RPE-1 data of changes in nuclear height are also included. Supplemental Figure 2 provides data for HT1080 FUCCI and MEF cell lines showing no change in actin contraction as measured by pMLC2 (phorphorylated myosin light chain 2) immunofluorescence throughout the interphase stages. Supplemental Figure 3 provides data in HT1080 cells showing decrease in H3K9me3 for larger vs. smaller nuclei and no change in lamin A/C.

## Acknowledgements

We would like to thank Dr. Kerry Bloom and Dr. Pierre Vidi for helpful and insightful discussions. We would like to thank lab members Kelsey Prince and Erin Walsh for helpful discussions. This work was primarily supported by NIH NIGMS grant Maximizing Investigators’ Research Award R35GM154928. This work has also been supported by the Center for 3D Structure and Physics of the Genome 4DN2 grant (1UM1 HG011536). The authors declare no competing interests.

## Data availability statement

The data are available in a public repository https://doi.org/10.6084/m9.figshare.28807298.v1

## Bibliography

Aureille, J, Buffière-Ribot, V, Harvey, BE, Boyault, C, Pernet, L, Andersen, T, Bacola, G, Balland, M, Fraboulet, S, Van Landeghem, L, et al. (2019). Nuclear envelope deformation controls cell cycle progression in response to mechanical force. EMBO Rep 20, e48084.

Banigan, EJ, Stephens, AD, and Marko, JF (2017). Mechanics and buckling of biopolymeric shells and cell nuclei. Biophys J 113, 1654–1663.

Belaghzal, H, Borrman, T, Stephens, AD, Lafontaine, DL, Venev, SV, Weng, Z, Marko, JF, and Dekker, J (2021). Liquid chromatin Hi-C characterizes compartment-dependent chromatin interaction dynamics. Nat Genet 53, 367–378.

Berg, IK, Currey, ML, Gupta, S, Berrada, Y, Nguyen, BV, Pho, M, Patteson, AE, Schwarz, JM, Banigan, EJ, and Stephens, AD (2023). Transcription inhibition suppresses nuclear blebbing and rupture independent of nuclear rigidity. J Cell Sci.

Bunner, S, Prince, K, Pujadas Liwag, EM, Eskndir, N, Srikrishna, K, McCarthy, AA, Kuklinski, A, Jackson, O, Pellegrino, P, Jagtap, S, et al. (2024). Decreased DNA density is a better indicator of a nuclear bleb than lamin B loss. J Cell Sci.

Camps, J, Wangsa, D, Falke, M, Brown, M, Case, CM, Erdos, MR, and Ried, T (2014). Loss of lamin B1 results in prolongation of S phase and decondensation of chromosome territories. FASEB J 28, 3423–3434.

Chakraborty, A, Prasanth, KV, and Prasanth, SG (2014). Dynamic phosphorylation of HP1α regulates mitotic progression in human cells. Nat Commun 5, 3445.

Chen, NY, Kim, P, Weston, TA, Edillo, L, Tu, Y, Fong, LG, and Young, SG (2018). Fibroblasts lacking nuclear lamins do not have nuclear blebs or protrusions but nevertheless have frequent nuclear membrane ruptures. Proc Natl Acad Sci U S A 115, 10100–10105.

Cho, S, Vashisth, M, Abbas, A, Majkut, S, Vogel, K, Xia, Y, Ivanovska, IL, Irianto, J, Tewari, M, Zhu, K, et al. (2019). Mechanosensing by the lamina protects against nuclear rupture, DNA damage, and cell-cycle arrest. Dev Cell 49, 920–935.e5.

Chu, CG, Lang, N, Walsh, E, Zheng, MD, Manning, G, Shalin, K, Cunha, LM, Faucon, KE, Kam, N, Folan, SN, et al. (2025). Lamin B loss in nuclear blebs is rupture dependent while increased DNA damage is rupture independent.

Currey, ML, Kandula, V, Biggs, R, Marko, JF, and Stephens, AD (2022). A versatile micromanipulation apparatus for biophysical assays of the cell nucleus. Cell Mol Bioeng.

Dahl, KN, Kahn, SM, Wilson, KL, and Discher, DE (2004). The nuclear envelope lamina network has elasticity and a compressibility limit suggestive of a molecular shock absorber. J Cell Sci 117, 4779–4786.

Danielsson, BE, Tieu, KV, Spagnol, ST, Vu, KK, Cabe, JI, Raisch, TB, Dahl, KN, and Conway, DE (2022). Chromatin condensation regulates endothelial cell adaptation to shear stress. Mol Biol Cell 33, ar101.

De Vos, WH, Houben, F, Kamps, M, Malhas, A, Verheyen, F, Cox, J, Manders, EMM, Verstraeten, VLRM, van Steensel, MAM, Marcelis, CLM, et al. (2011). Repetitive disruptions of the nuclear envelope invoke temporary loss of cellular compartmentalization in laminopathies. Hum Mol Genet 20, 4175–4186.

Dechat, T, Gajewski, A, Korbei, B, Gerlich, D, Daigle, N, Haraguchi, T, Furukawa, K, Ellenberg, J, and Foisner, R (2004). LAP2alpha and BAF transiently localize to telomeres and specific regions on chromatin during nuclear assembly. J Cell Sci 117, 6117–6128.

Denais, CM, Gilbert, RM, Isermann, P, McGregor, AL, te Lindert, M, Weigelin, B, Davidson, PM, Friedl, P, Wolf, K, and Lammerding, J (2016). Nuclear envelope rupture and repair during cancer cell migration. Science 352, 353–358.

Earle, AJ, Kirby, TJ, Fedorchak, GR, Isermann, P, Patel, J, Iruvanti, S, Moore, SA, Bonne, G, Wallrath, LL, and Lammerding, J (2020). Mutant lamins cause nuclear envelope rupture and DNA damage in skeletal muscle cells. Nat Mater 19, 464–473.

Furusawa, T, Rochman, M, Taher, L, Dimitriadis, EK, Nagashima, K, Anderson, S, and Bustin, M (2015). Chromatin decompaction by the nucleosomal binding protein HMGN5 impairs nuclear sturdiness. Nat Commun 6, 6138.

Gratzner, HG, Leif, RC, Ingram, DJ, and Castro, A (1975). The use of antibody specific for bromodeoxyuridine for the immunofluorescent determination of DNA replication in single cells and chromosomes. Exp Cell Res 95, 88–94.

Greiner, D, Bonaldi, T, Eskeland, R, Roemer, E, and Imhof, A (2005). Identification of a specific inhibitor of the histone methyltransferase SU(VAR)3-9. Nat Chem Biol 1, 143–145.

Gurvich, N, Tsygankova, OM, Meinkoth, JL, and Klein, PS (2004). Histone deacetylase is a target of valproic acid-mediated cellular differentiation. Cancer Res 64, 1079–1086.

Harr, JC, Luperchio, TR, Wong, X, Cohen, E, Wheelan, SJ, and Reddy, KL (2015). Directed targeting of chromatin to the nuclear lamina is mediated by chromatin state and A-type lamins. J Cell Biol 208, 33–52.

Hatch, EM, and Hetzer, MW (2016). Nuclear envelope rupture is induced by actin-based nucleus confinement. J Cell Biol 215, 27–36.

Helfand, BT, Wang, Y, Pfleghaar, K, Shimi, T, Taimen, P, and Shumaker, DK (2012). Chromosomal regions associated with prostate cancer risk localize to lamin B-deficient microdomains and exhibit reduced gene transcription. J Pathol 226, 735–745.

Hobson, CM, Kern, M, O’Brien, ET, 3rd, Stephens, AD, Falvo, MR, and Superfine, R (2020). Correlating nuclear morphology and external force with combined atomic force microscopy and light sheet imaging separates roles of chromatin and lamin A/C in nuclear mechanics. Mol Biol Cell 31, 1788–1801.

Iida, S, Shinkai, S, Itoh, Y, Tamura, S, Kanemaki, MT, Onami, S, and Maeshima, K (2022). Single-nucleosome imaging reveals steady-state motion of interphase chromatin in living human cells. Sci Adv 8, eabn5626.

Irianto, J, Pfeifer, CR, Bennett, RR, Xia, Y, Ivanovska, IL, Liu, AJ, Greenberg, RA, and Discher, DE (2016). Nuclear constriction segregates mobile nuclear proteins away from chromatin. Mol Biol Cell 27, 4011–4020.

Irianto, J, Xia, Y, Pfeifer, CR, Athirasala, A, Ji, J, Alvey, C, Tewari, M, Bennett, RR, Harding, SM, Liu, AJ, et al. (2017). DNA damage follows repair factor depletion and portends genome variation in cancer cells after pore migration. Curr Biol 27, 210–223.

Jones, MC, Askari, JA, Humphries, JD, and Humphries, MJ (2018). Cell adhesion is regulated by CDK1 during the cell cycle. J Cell Biol 217, 3203–3218.

Kalukula, Y, Stephens, AD, Lammerding, J, and Gabriele, S (2022). Mechanics and functional consequences of nuclear deformations. Nat Rev Mol Cell Biol.

Kamikawa, Y, Wu, Z, Nakazawa, N, Ito, T, Saito, A, and Imaizumi, K (2023). Impact of cell cycle on repair of ruptured nuclear envelope and sensitivity to nuclear envelope stress in glioblastoma. Cell Death Discov 9, 233.

Kim, D-H, Khatau, SB, Feng, Y, Walcott, S, Sun, SX, Longmore, GD, and Wirtz, D (2012). Actin cap associated focal adhesions and their distinct role in cellular mechanosensing. Sci Rep 2, 555.

Lammerding, J, Fong, LG, Ji, JY, Reue, K, Stewart, CL, Young, SG, and Lee, RT (2006). Lamins A and C but not lamin B1 regulate nuclear mechanics. J Biol Chem 281, 25768–25780.

Le Berre, M, Aubertin, J, and Piel, M (2012). Fine control of nuclear confinement identifies a threshold deformation leading to lamina rupture and induction of specific genes. Integr Biol (Camb) 4, 1406–1414.

Manning, G, Li, A, Eskndir, N, Currey, M, and Stephens, AD (2025). Constitutive heterochromatin controls nuclear mechanics, morphology, and integrity through H3K9me3 mediated chromocenter compaction. Nucleus 16.

Marcus, JM, Burke, RT, DeSisto, JA, Landesman, Y, and Orth, JD (2015). Longitudinal tracking of single live cancer cells to understand cell cycle effects of the nuclear export inhibitor, selinexor. Sci Rep 5, 14391.

Miranda, TB, Cortez, CC, Yoo, CB, Liang, G, Abe, M, Kelly, TK, Marquez, VE, and Jones, PA (2009). DZNep is a global histone methylation inhibitor that reactivates developmental genes not silenced by DNA methylation. Mol Cancer Ther 8, 1579–1588.

Mistriotis, P, Wisniewski, EO, Bera, K, Keys, J, Li, Y, Tuntithavornwat, S, Law, RA, Perez-Gonzalez, NA, Erdogmus, E, Zhang, Y, et al. (2019). Confinement hinders motility by inducing RhoA-mediated nuclear influx, volume expansion, and blebbing. J Cell Biol 218, 4093–4111.

Nader, GP de F, Agüera-Gonzalez, S, Routet, F, Gratia, M, Maurin, M, Cancila, V, Cadart, C, Palamidessi, A, Ramos, RN, San Roman, M, et al. (2021). Compromised nuclear envelope integrity drives TREX1-dependent DNA damage and tumor cell invasion. Cell 184, 5230–5246.e22.

Nagano, T, Lubling, Y, Varnai, C, Dudley, C, Leung, W, Baran, Y, Mandelson-Cohen, N, Wingett, S, Fraser, P, and Tanay, A (2016). Cell cycle dynamics of chromosomal organisation at single-cell resolution, bioRxiv.

Nava, MM, Miroshnikova, YA, Biggs, LC, Whitefield, DB, Metge, F, Boucas, J, Vihinen, H, Jokitalo, E, Li, X, García Arcos, JM, et al. (2020). Heterochromatin-driven nuclear softening protects the genome against mechanical stress-induced damage. Cell 181, 800–817.e22.

Nmezi, B, Xu, J, Fu, R, Armiger, TJ, Rodriguez-Bey, G, Powell, JS, Ma, H, Sullivan, M, Tu, Y, Chen, NY, et al. (2019). Concentric organization of A- and B-type lamins predicts their distinct roles in the spatial organization and stability of the nuclear lamina. Proc Natl Acad Sci U S A 116, 4307–4315.

Pfeifer, CR, Xia, Y, Zhu, K, Liu, D, Irianto, J, García, VMM, Millán, LMS, Niese, B, Harding, S, Deviri, D, et al. (2018). Constricted migration increases DNA damage and independently represses cell cycle. Mol Biol Cell 29, 1948–1962.

Pho, M, Berrada, Y, Gunda, A, Lavallee, A, Chiu, K, Padam, A, Currey, ML, and Stephens, AD (2023). Actin contraction controls nuclear blebbing and rupture independent of actin confinement. Mol Biol Cell, mbcE23070292.

Pujadas Liwag, EM, Acosta, N, Almassalha, LM, Su, YP, Gong, R, Kanemaki, MT, Stephens, AD, and Backman, V (2025). Nuclear blebs are associated with destabilized chromatin-packing domains. J Cell Sci 138.

Raab, M, Gentili, M, de Belly, H, Thiam, H-R, Vargas, P, Jimenez, AJ, Lautenschlaeger, F, Voituriez, R, Lennon-Duménil, A-M, Manel, N, et al. (2016). ESCRT III repairs nuclear envelope ruptures during cell migration to limit DNA damage and cell death. Science 352, 359–362.

Rao, S, Porter, DC, Chen, X, Herliczek, T, Lowe, M, and Keyomarsi, K (1999). Lovastatin-mediated G1 arrest is through inhibition of the proteasome, independent of hydroxymethyl glutaryl-CoA reductase. Proc Natl Acad Sci U S A 96, 7797–7802.

Sakaue-Sawano, A, Kurokawa, H, Morimura, T, Hanyu, A, Hama, H, Osawa, H, Kashiwagi, S, Fukami, K, Miyata, T, Miyoshi, H, et al. (2008). Visualizing spatiotemporal dynamics of multicellular cell-cycle progression. Cell 132, 487–498.

van Schaik, T, Vos, M, Peric-Hupkes, D, Hn Celie, P, and van Steensel, B (2020). Cell cycle dynamics of lamina-associated DNA. EMBO Rep 21, e50636.

Senigagliesi, B, Penzo, C, Severino, LU, Maraspini, R, Petrosino, S, Morales-Navarrete, H, Pobega, E, Ambrosetti, E, Parisse, P, Pegoraro, S, et al. (2019). The High Mobility Group A1 (HMGA1) chromatin architectural factor modulates nuclear stiffness in breast cancer cells. Int J Mol Sci 20, 2733.

Shah, P, Hobson, CM, Cheng, S, Colville, MJ, Paszek, MJ, Superfine, R, and Lammerding, J (2021). Nuclear deformation causes DNA damage by increasing replication stress. Curr Biol 31, 753–765.e6.

Shimamoto, Y, Tamura, S, Masumoto, H, and Maeshima, K (2017). Nucleosome-nucleosome interactions via histone tails and linker DNA regulate nuclear rigidity. Mol Biol Cell 28, 1580– 1589.

Stephens, AD, Banigan, EJ, Adam, SA, Goldman, RD, and Marko, JF (2017). Chromatin and lamin A determine two different mechanical response regimes of the cell nucleus. Mol Biol Cell 28, 1984–1996.

Stephens, AD, Banigan, EJ, and Marko, JF (2019a). Chromatin’s physical properties shape the nucleus and its functions. Curr Opin Cell Biol 58, 76–84.

Stephens, AD, Liu, PZ, Banigan, EJ, Almassalha, LM, Backman, V, Adam, SA, Goldman, RD, and Marko, JF (2018). Chromatin histone modifications and rigidity affect nuclear morphology independent of lamins. Mol Biol Cell 29, 220–233.

Stephens, AD, Liu, PZ, Kandula, V, Chen, H, Almassalha, LM, Herman, C, Backman, V, O’Halloran, T, Adam, SA, Goldman, RD, et al. (2019b). Physicochemical mechanotransduction alters nuclear shape and mechanics via heterochromatin formation. Mol Biol Cell 30, 2320– 2330.

Strom, AR, Biggs, RJ, Banigan, EJ, Wang, X, Chiu, K, Herman, C, Collado, J, Yue, F, Ritland Politz, JC, Tait, LJ, et al. (2021). HP1α is a chromatin crosslinker that controls nuclear and mitotic chromosome mechanics. Elife 10.

Swift, J, Ivanovska, IL, Buxboim, A, Harada, T, Dingal, PCDP, Pinter, J, Pajerowski, JD, Spinler, KR, Shin, J-W, Tewari, M, et al. (2013). Nuclear lamin-A scales with tissue stiffness and enhances matrix-directed differentiation. Science 341, 1240104.

Vahabikashi, A, Sivagurunathan, S, Nicdao, FAS, Han, YL, Park, CY, Kittisopikul, M, Wong, X, Tran, JR, Gundersen, GG, Reddy, KL, et al. (2022). Nuclear lamin isoforms differentially contribute to LINC complex-dependent nucleocytoskeletal coupling and whole-cell mechanics. Proc Natl Acad Sci U S A 119, e2121816119.

Vassilev, LT, Tovar, C, Chen, S, Knezevic, D, Zhao, X, Sun, H, Heimbrook, DC, and Chen, L (2006). Selective small-molecule inhibitor reveals critical mitotic functions of human CDK1. Proc Natl Acad Sci U S A 103, 10660–10665.

Vianay, B, Senger, F, Alamos, S, Anjur-Dietrich, M, Bearce, E, Cheeseman, B, Lee, L, and Théry, M (2018). Variation in traction forces during cell cycle progression. Biol Cell 110, 91–96.

Williams, JF, Surovtsev, IV, Schreiner, SM, Chen, Z, Raiymbek, G, Nguyen, H, Hu, Y, Biteen, JS, Mochrie, SGJ, Ragunathan, K, et al. (2024). The condensation of HP1α/Swi6 imparts nuclear stiffness. Cell Rep 43, 114373.

Williams, JF, Surovtsev, IV, Schreiner, SM, Nguyen, H, Hu, Y, Mochrie, SGJ, and King, MC (2020). Phase separation enables heterochromatin domains to do mechanical work, bioRxiv.

Xia, Y, Ivanovska, IL, Zhu, K, Smith, L, Irianto, J, Pfeifer, CR, Alvey, CM, Ji, J, Liu, D, Cho, S, et al. (2018). Nuclear rupture at sites of high curvature compromises retention of DNA repair factors. J Cell Biol 217, 3796–3808.

Xia, Y, Pfeifer, CR, Zhu, K, Irianto, J, Liu, D, Pannell, K, Chen, EJ, Dooling, LJ, Tobin, MP, Wang, M, et al. (2019). Rescue of DNA damage after constricted migration reveals a mechano-regulated threshold for cell cycle. J Cell Biol 218, 2545–2563.

