## Supplemental Figures 1-3 for "Changes in nuclear and actin mechanics from G1 to G2 affect nuclear integrity"

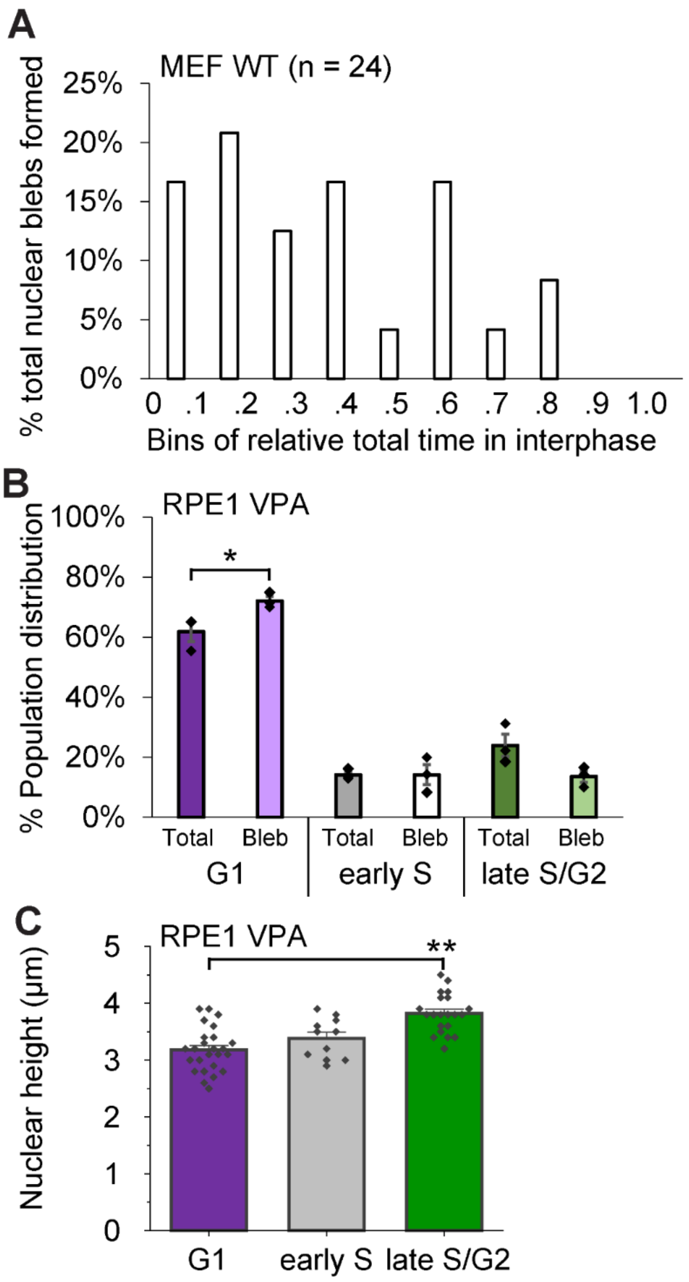


**Supplemental Figure 1. Increased bleb formation in G1 occurs across cell lines.** (A) Graph of MEF wild type (WT) NLS-GFP cells percentage of all nuclear blebs formed relative to that cell's normalized interphase cell cycle length (n = 24, G1 67%, early S 25%, late S/G2 8 %). Cells were tracked for 36 hours at 30-minute intervals to allow for watch the same cell end of the first mitosis through interphase till the start of the next mitosis. (B) Graph of RPE1 FUCCI VPA-treated cells percentage of all cells for all (total) compared to percentage of all nuclear bleb formation events (bleb) in G1, early S, and late S/G2. Biological triplicate n = 29,56,79. (C) Graph of nuclear height for RPE1 FUCCI VPA-treated cells for G1, early S, and late S/G2 (n = 25,11,21 respectively). Unpaired two-tailed Student's t-test p values reported as *<0.05, **<0.01, ***<0.001, or ns denotes no significance, p>0.05. Error bars represent standard error.

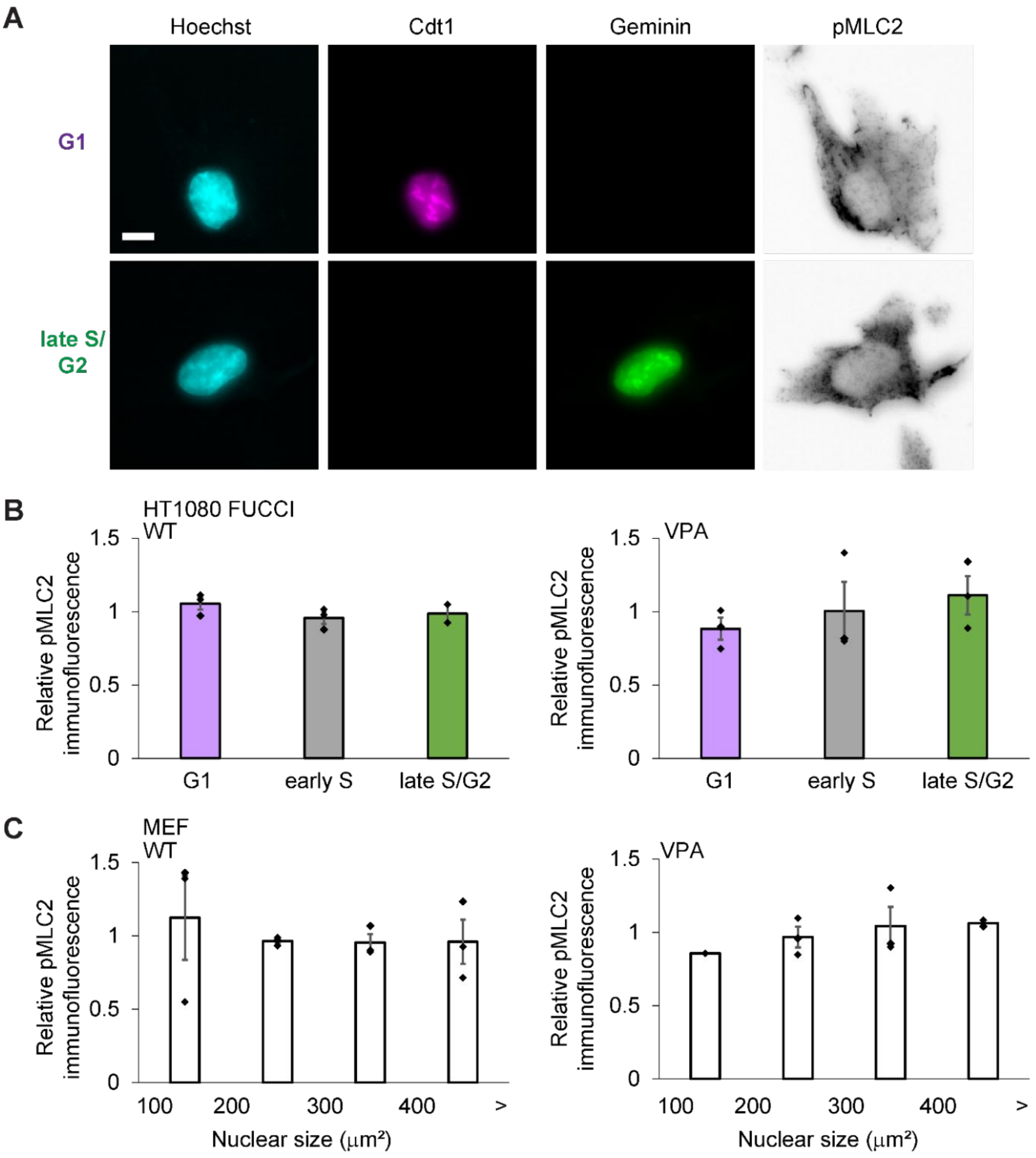


**Supplemental Figure 2. Actin contraction levels are similar across the interphase stages of the cell cycle.** (A) Example images of a G1 and late S/G2 nucleus stained for DNA via Hoechst, containing fluorescent Cdt1and Geminin, and immunofluorescence for phosphorylated myosin light chain 2 (pMLC2). (B) Graph of relative pMLC2 immunofluorescence for G1, early S, and late S/G2 in HT1080 FUCCI wild type (WT) and chromatin decompaction (VPA). (C) Graph of relative pMLC2 immunofluorescence for different nuclear sizes as a proxy for interphase stage in MEF wild type (WT) and chromatin decompaction treatment (VPA). Biological triplicate with n > 20 cells each condition. Unpaired two-tailed Student's t-test p values where all p>0.05, no significance. Error bars represent standard error. Scale bar = 10 µm.

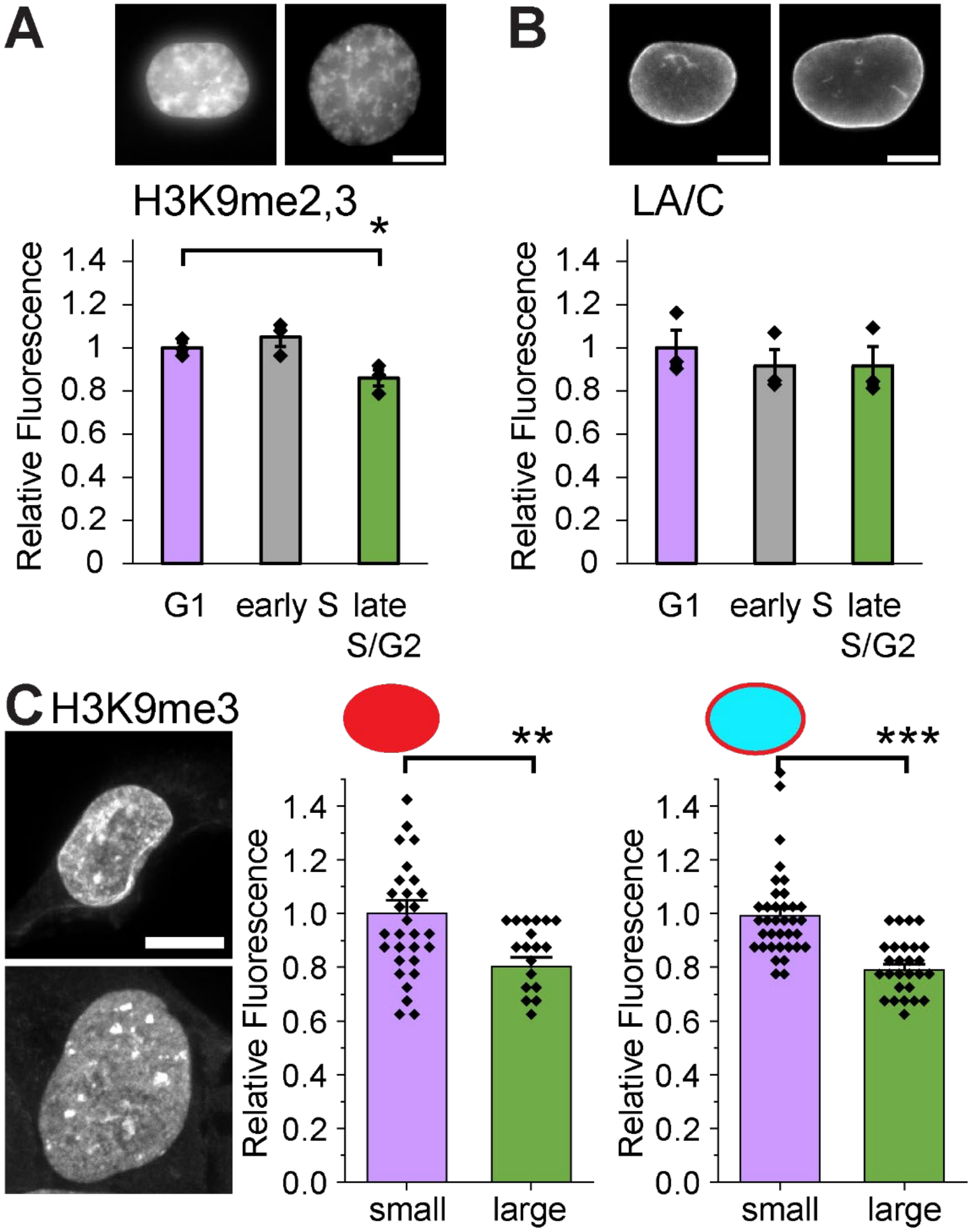


**Supplemental Figure 3. Constitutive heterochromatin is decreased from G1 to late S/G2 while lamin A/C remains similar.** (A,B) Example images and graphs of HT1080 FUCCI G1, early S, and late S/G2 relative immunofluorescence intensity for (A) constitutive heterochromatin marker H3K9me2,3 and (B) lamin A/C (N = 3 where each contained > 20 nuclei for each condition). (C) HT1080 confocal imaging of H3K9me3 example images and graphs of whole (left) and periphery (right) nucleus relative fluorescence intensity for small G1-like vs larger G2-like nuclei. Unpaired two-tailed Student's t-test p values reported as *<0.05, **<0.01, ***<0.001, or ns denotes no significance, p>0.05. Error bars represent standard error. Scale bar = 10 µm.

**Video 1.** HT1080 FUCCI cells imaged for nuclear bleb formation throughout the interphase cell cycle.

**Video 2.** HT1080 FUCCI cells imaged while being placed under artificial confinement to 4 μm.
